# The E3/E4 ubiquitin conjugation factor UBE4B interacts with and ubiquitinates the HTLV-1 Tax oncoprotein to promote NF-κB activation

**DOI:** 10.1101/2020.03.30.016253

**Authors:** Suchitra Mohanty, Teng Han, Young Bong Choi, Alfonso Lavorgna, Jiawen Zhang, Edward W. Harhaj

**Author notes:** Equal contribution (Suchitra Mohanty, Teng Han). Weill Cornell Medicine, New York, NY 10021. MilliporeSigma, Rockville, MD 20850.

## Abstract

Human T-cell leukemia virus type 1 (HTLV-1) is the etiological agent of adult T-cell leukemia/lymphoma (ATLL), and the neurological disease HTLV-1 associated myelopathy/tropical spastic paraparesis (HAM/TSP). The HTLV-1 Tax protein persistently activates the NF-κB pathway to enhance the proliferation and survival of HTLV-1 infected T cells. Lysine 63 (K63)-linked polyubiquitination of Tax provides an important regulatory mechanism that promotes Tax-mediated interaction with the IKK complex and activation of NF-κB; however, the host proteins regulating Tax ubiquitination are largely unknown. To identify novel Tax interacting proteins that may regulate its ubiquitination we conducted a yeast two-hybrid screen using Tax as bait. This screen yielded the E3/E4 ubiquitin conjugation factor UBE4B as a novel binding partner for Tax. Here, we confirmed the interaction between Tax and UBE4B in mammalian cells by co-immunoprecipitation assays and demonstrated colocalization by proximity ligation assay and confocal microscopy. Overexpression of UBE4B specifically enhanced Tax-induced NF-κB activation, whereas knockdown of UBE4B impaired Tax-induced NF-κB activation and the induction of NF-κB target genes in T cells and ATLL cell lines. Furthermore, depletion of UBE4B with shRNA resulted in apoptotic cell death and diminished the proliferation of ATLL cell lines. Finally, overexpression of UBE4B enhanced Tax polyubiquitination, and knockdown or CRISPR/Cas9-mediated knockout of UBE4B attenuated both K48- and K63-linked polyubiquitination of Tax. Collectively, these results implicate UBE4B in HTLV-1 Tax polyubiquitination and downstream NF-κB activation.

**Author Summary:** Infection with the retrovirus HTLV-1 leads to the development of either CD4+CD25+ leukemia/lymphoma (ATLL) or a demyelinating neuroinflammatory disease (HAM/TSP) in a subset of infected individuals. The HTLV-1 Tax protein is a regulatory protein which regulates viral gene expression and persistently activates cellular signaling pathways such as NF-κB to drive the clonal expansion and longevity of HTLV-1 infected CD4+ T cells. Polyubiquitination of Tax is a key mechanism of NF-κB activation by assembling and activating IκB kinase (IKK) signaling complexes; however, the host factors regulating Tax ubiquitination have remained elusive. Here, we have identified the E3/E4 ubiquitin conjugation factor UBE4B as a novel Tax binding protein that promotes both K48- and K63-linked polyubiquitination of Tax. Knockdown or knockout of UBE4B impairs Tax-induced NF-κB activation and triggers apoptosis of HTLV-1 transformed cells. Therefore, UBE4B is an integral host factor that supports HTLV-1 Tax polyubiquitination, NF-κB activation and cell survival.

## Introduction

Human T-cell leukemia virus type 1 (HTLV-1) is an oncogenic retrovirus estimated to infect between 5 and 10 million people worldwide [1]. Highly endemic areas of HTLV-1 include southern Japan, sub-Saharan Africa, South America, the Caribbean and the Middle East. Recent epidemiological studies have found >40% of adults in indigenous communities in Central Australia infected with HTLV-1 [2]. HTLV-1 predominantly infects CD4+ T cells and infection is associated with the development of a CD4+CD25+ malignancy, adult T-cell leukemia/lymphoma (ATLL), in ∼5% of infected individuals after a long latent period (about 60 years) [3]. HTLV-1 infection can also lead to inflammatory diseases such as HTLV-1-associated myelopathy/tropical spastic paraparesis (HAM/TSP) [4].

The HTLV-1 regulatory proteins Tax and HBZ are strongly linked to viral persistence and pathogenesis [5]. Both Tax and HBZ have been ascribed oncogenic functions and together contribute to the survival of HTLV-1 infected cells, and play distinct roles in the genesis and/or maintenance of ATLL. Tax is required for viral gene expression by recruiting CREB/ATF transcription factors and coactivators CBP/p300 to the HTLV-1 promoter [6]. In addition, Tax dysregulates the cell cycle, inhibits tumor suppressors Rb and p53, and activates multiple signaling pathways such as NF-κB to enforce a transcriptional program supporting the proliferation and survival of T cells carrying the HTLV-1 provirus [7]. Tax is highly immunogenic and represents a main target of CD8+ T cells, therefore Tax expression and HTLV-1 plus-strand transcription is commonly silenced as a mechanism of immune evasion [8]. However, the HTLV-1 provirus can be reactivated from latency in response to stress stimuli such as hypoxia, p38-MAPK signaling or oxidative stress [9–11]. In ATLL, Tax expression is lost in ∼60% of tumors due to genetic or epigenetic mechanisms [12]; however, HBZ expression remains ubiquitously expressed.

Tax is a potent activator of canonical and noncanonical NF-κB pathways, and persistent activation of NF-κB is a hallmark of ATLL [13]. Tax interacts with the IκB kinase (IKK) complex through direct binding with NEMO/IKKγ to promote the phosphorylation and degradation of the inhibitor IκBα [14]. Modification of Tax by polyubiquitination is tightly linked to NF-κB activation. Tax is conjugated with lysine 63 (K63)-linked polyubiquitin (polyUb) chains, resulting in the recruitment of the IKK complex to foci in the vicinity of the cis-Golgi, which serves as a hub for Tax/IKK interaction and IKK activation [15]. Tax K63-linked polyubiquitination on multiple C-terminal lysines facilitates interaction with NEMO and the IKK complex for the subsequent activation of NF-κB [16]. The E2 enzyme Ubc13 plays a critical role in the K63-linked polyubiquitination of Tax and NF-κB activation [17]; however, other host factors directly modulating Tax ubiquitination are largely unknown. The E3 ligase PDLIM2 has been shown to conjugate K48-linked polyUb chains on Tax, however this leads to the proteasomal degradation of Tax in the nuclear matrix [18]. Tax also interacts with selective autophagy receptors TAX1BP1, Optineurin and SQSTM-1/p62, to potentiate Tax ubiquitination and/or NF-κB activation [19–21], although the role of autophagy in Tax-IKK activation remains poorly understood. Finally, Tax can interact with membrane associated CADM1/TSLC1 to promote K63-linked polyubiquitination of Tax and the inhibition of the ubiquitin-editing enzyme A20 [22].

Tax has also been shown to target E3 ligases for the generation of unanchored polyUb chains or ubiquitination of substrates other than Tax to activate IKK signaling or promote cell survival. Tax activates Ring Finger Protein 8 (RNF8) to induce the assembly of long unanchored K63-linked polyUb chains that activate IKK and TAK1 kinases [23]. Tax has also been shown to recruit the linear (Met1-linked) ubiquitin chain assembly complex (LUBAC) to IKK for the generation of K63/M1-linked hybrid polyUb chains [24]. These hybrid chains are thought to trigger the oligomerization and activation of the IKK complex. Tax interacts with the E3 ligase TRAF6 and promotes its E3 activity, leading to MCL-1 K63-linked polyubiquitination and stabilization and inhibition of apoptosis [25]. Although Tax binds to TRAF6 it is unclear if TRAF6 can conjugate Tax with K63-linked polyUb chains. Taken together, ubiquitination plays substantial roles in Tax-induced IKK activation and oncogenesis.

UBE4B is a human homolog of the *Saccharomyces cerevisiae* UFD2. Yeast UFD2, encoded by a single-copy gene, is the first reported E4 ubiquitination factor and is required for elongation of an oligoubiquitin chain on certain substrates that are subsequently recognized by the 26S proteasome for degradation [26, 27]. UFD2 is involved in the endoplasmic reticulum-associated degradation (ERAD) pathway by interacting with the AAA-type ATPase Cdc48 [28]. UBE4B and its homologs share a 70-amino acid U-box domain that confers ligase activity. Sequence profile alignments indicate that the U-box is a derived version of the RING finger domain, but the signature cysteines in the RING finger domain that are responsible for metal chelating, are not conserved in the U-box [29]. However, the predicted structure of the U-box is very similar to that of the RING finger domain [30], which indicates that U-box proteins may also have the capability to function as E3 ligases independently. Indeed, UFD2 can function as a bona fide E3 ubiquitin ligase to promote ubiquitin conjugation of unfolded proteins [31]. Thus, UBE4B and its homologs can clearly function as E3 ligases, and their E4 function may represent a specialized type of E3 activity with mono- or oligoubiquitinated proteins as substrates. UBE4B has been identified as an E3/E4 ligase that collaborates with Hdm2 to catalyze the polyubiquitination and degradation of the tumor suppressor p53 [32]. UBE4B has also been shown to ubiquitinate additional substrates such as EGFR, Ataxin-3 and OTUB1 [33–35]. To date, UBE4B has not been implicated in the ubiquitination of any viral proteins.

In this study, we undertook an approach to identify novel Tax binding proteins that may regulate Tax polyubiquitination and NF-κB activation. We have identified the E3/E4 ubiquitin conjugation factor UBE4B as a novel Tax binding protein which promotes both K63- and K48-linked Tax polyubiquitination and NF-κB activation.

## Results

### Tax interacts with UBE4B

In order to identify new binding partners of Tax, we conducted a yeast two-hybrid screen using full-length Tax as bait with a cDNA library derived from human leukocytes and activated mononuclear cells. Importantly, known Tax interacting proteins DLG1 [36], NF-κB2 [37] and TAX1BP3 [38] were identified in the screen. A potential novel Tax binding protein that emerged from this screen was the E3/E4 ubiquitin conjugation factor UBE4B. To confirm the interaction between Tax and UBE4B, a co-immunoprecipitation (co-IP) experiment was performed with lysates from 293T cells transfected with Tax and a Flag-tagged UBE4B plasmid. As shown in Figure 1A, UBE4B interacted with Tax when both proteins were overexpressed. The reciprocal IP in which UBE4B was immunoprecipitated also confirmed Tax and UBE4B interaction (Figure 1B). The interaction was further examined by co-IPs using Tax and a UBE4B catalytically inactive mutant, in which the highly conserved proline at position 1140 was replaced with alanine (UBE4B P1140A) [39]. The dominant-negative mutant (UBE4B P1140A) was still able to interact with Tax (Figure S1A). Tax point mutants M22 and M47 are impaired in NF-κB and CREB activation, respectively [17]. UBE4B interacted with Tax M22 and M47 mutants, similar to Tax WT (Figure S1B).

**Figure 1.**
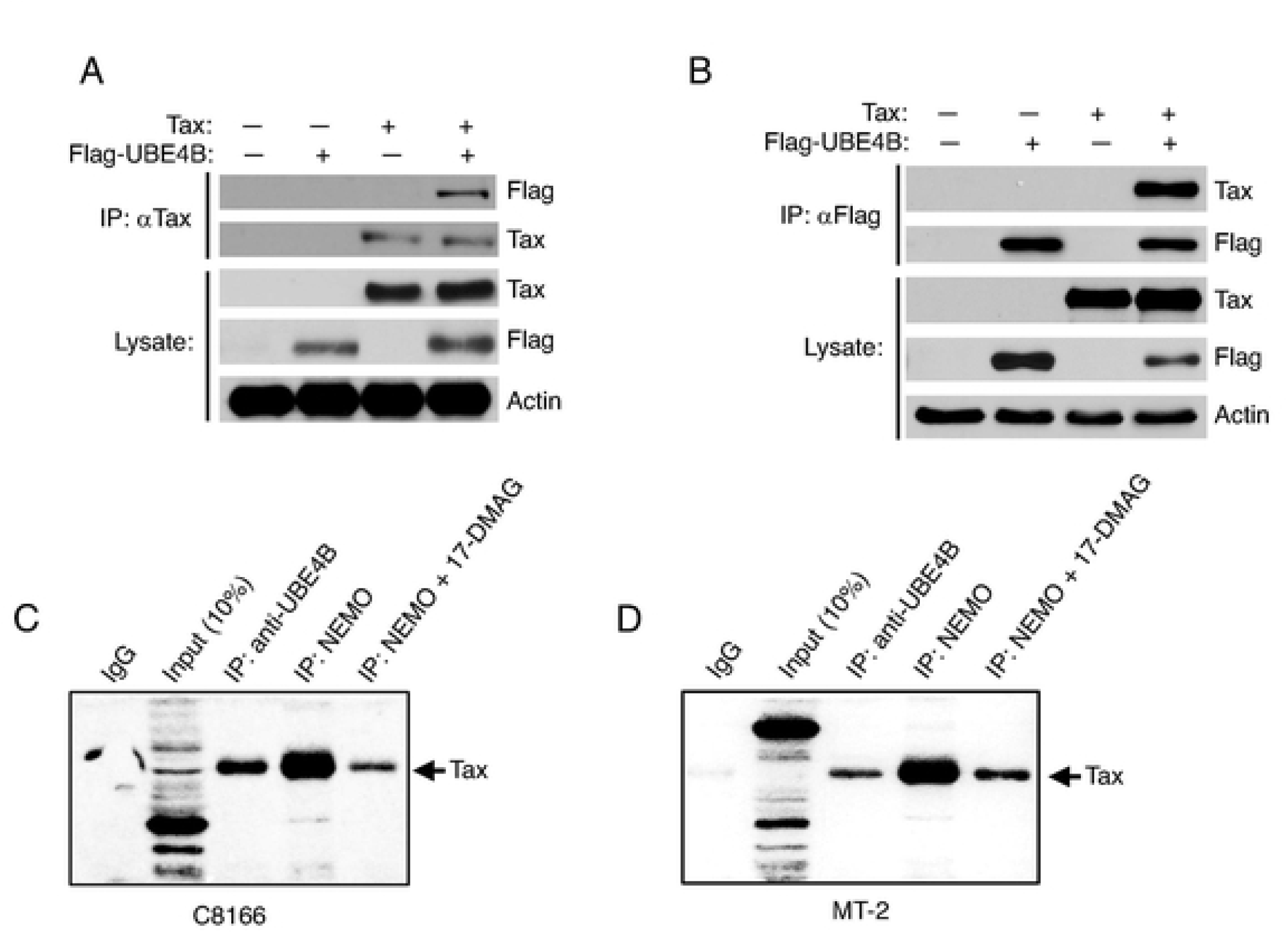
Tax interacts with UBE4B. (A, B) Co-IP analysis with either Tax (A) or Flag-UBE4B (B) immunoprecipitates from lysates of 293T cells transfected with the indicated plasmids. (C, D) Co-IP analysis with either control IgG, anti-UBE4B or anti-NEMO immunoprecipitates from lysates of C8166 and MT-2 cells either untreated or treated with 17-DMAG as indicated.

We next performed co-IPs using HTLV-1 transformed cell lines C8166 and MT-2 to determine if endogenous Tax and UBE4B proteins could interact. The anti-UBE4B-immunoprecipitated complex was immunoblotted with anti-Tax. As a positive control, NEMO was immunoprecipitated and immunoblotted with anti-Tax. As expected, NEMO and Tax strongly interacted in both C8166 and MT-2 cells (Figure 1C, D). UBE4B was also found to interact with Tax using lysates from C8166 and MT-2 cells (Figure 1C, D). To confirm the specificity of the Tax immunoreactive band in the co-IPs, cells were treated with the heat shock protein 90 (HSP90) inhibitor 17-DMAG which induces Tax degradation [40]. Treatment of C8166 and MT-2 cells with 17-DMAG reduced the amount of Tax that was immunoprecipitated with NEMO (Figure 1C, D). Therefore, UBE4B interacts with Tax under both overexpression and endogenous conditions.

### UBE4B and Tax colocalize in HTLV-1 transformed cells

Tax shuttles between the cytoplasm and nucleus and is distributed in both compartments at steady state [41]. Similarly, UBE4B localizes in both the cytoplasm and nucleus [33], therefore we sought to determine where Tax and UBE4B interacted in cells. First, we fractionated MT-2, HUT-102 and C8166 cells into cytoplasmic and nuclear fractions, and examined UBE4B and Tax expression. Lactate dehydrogenase (LDH) and poly (ADP-ribose) polymerase (PARP) immunoblotting confirmed the purity of cytoplasmic and nuclear fractions, respectively. As expected, Tax was expressed in both cytoplasmic and nuclear compartments in the cell lines (Figure 2A). UBE4B was found predominantly in cytoplasmic fractions, but was also detected in nuclear fractions (Figure 2A). Next, MT-2 and C8166 cells were subjected to immunostaining and confocal microscopy. Tax strongly co-localized with UBE4B, predominantly in the cytoplasm, in both MT-2 and C8166 cells (Figure 2B). To further confirm the Tax-UBE4B interaction, we conducted an in situ proximity ligation assay (PLA), which can detect proteins found within 40 nm of each other. PLA signals were only observed in MT-2 and C8166 cells when both Tax and UBE4B antibodies were used together with secondary probes (Figure 2C). PLA signals were found mostly in the cytoplasm, but were also detected in the nucleus. PLA signals were quantified, confirming a significant increase compared to secondary probe alone (Figure 2D). These data provide further evidence of an interaction between endogenous Tax and UBE4B.

**Figure 2.**
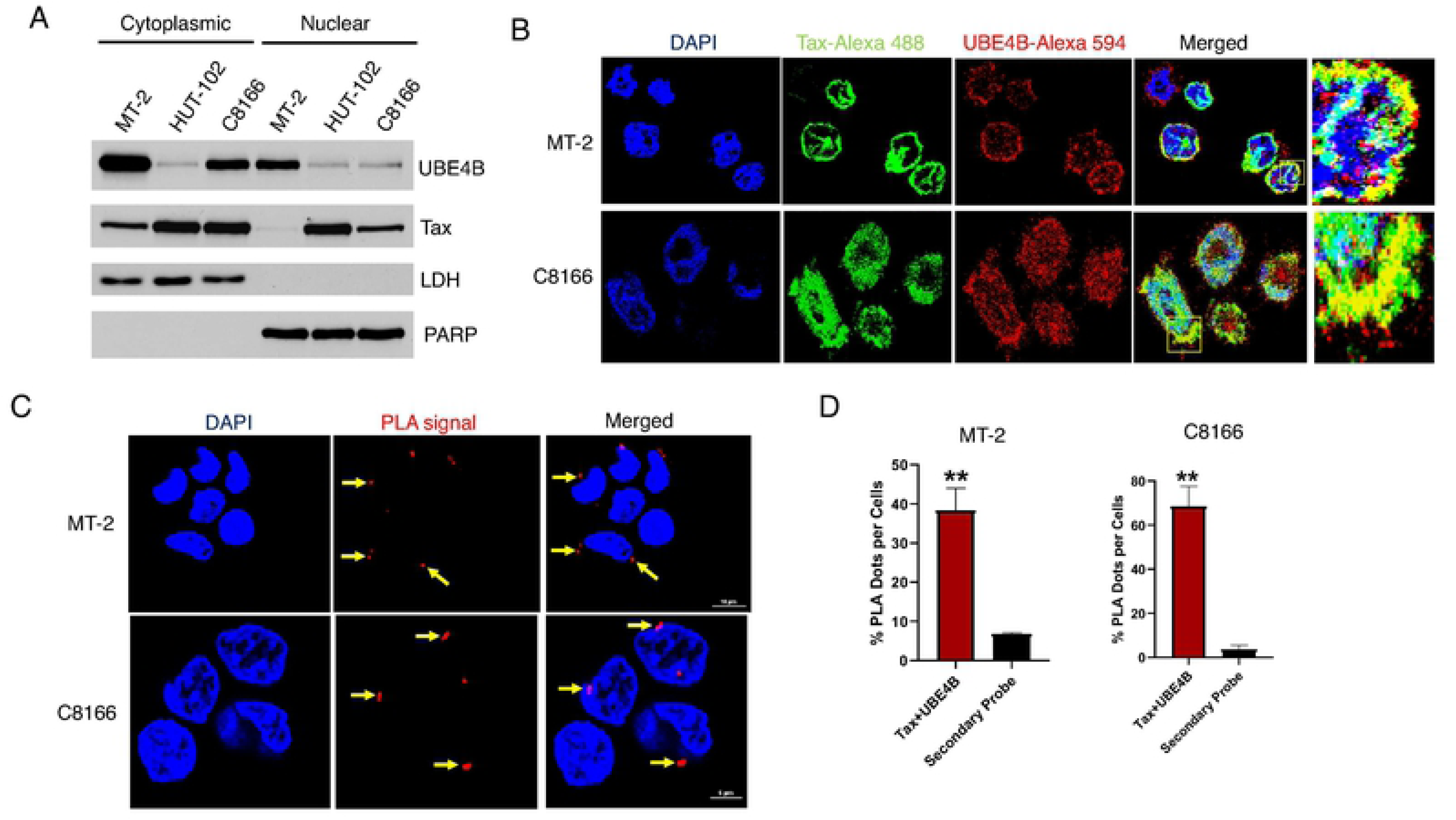
Co-localization of UBE4B and Tax in HTLV-1 transformed cells. (A) Immunoblotting was performed with the indicated antibodies using cytoplasmic and nuclear fractions obtained from MT-2, HUT-102 and C8166 cells. (B) Immunofluorescence (IF) confocal microscopy was performed using MT-2 and C8166 cells with the indicated antibodies. (C) PLA was performed using MT-2 and C8166 cells with Tax or UBE4B antibodies. (D) Graphical representation of the number of PLA signals with the secondary probe stained sample as a negative control.

### UBE4B enhances Tax-mediated activation of NF-κB

Since UBE4B and Tax mainly interacted in the cytoplasm, where Tax activates IKK, we hypothesized that UBE4B may regulate Tax-induced NF-κB activation. To test this hypothesis, 293T cells were transfected with Flag-Tax together with Flag-UBE4B or catalytically inactive Flag-UBE4B P1140A, and NF-κB luciferase assays were performed. As expected, Tax expression resulted in potent activation of the NF-κB luciferase reporter (Figure 3A). Transfection of wild-type UBE4B, but not catalytically inactive UBE4B P1140A, significantly enhanced Tax-mediated NF-κB activation (Figure 3A). Wild-type UBE4B or UBE4B P1140A alone had no effect on NF-κB activation (Figure 3A). In contrast, overexpression of UBE4B had no effect on Tax-mediated HTLV-1 LTR activation, therefore UBE4B appears to selectively modulate Tax activation of NF-κB (Figure 3B).

**Figure 3.**
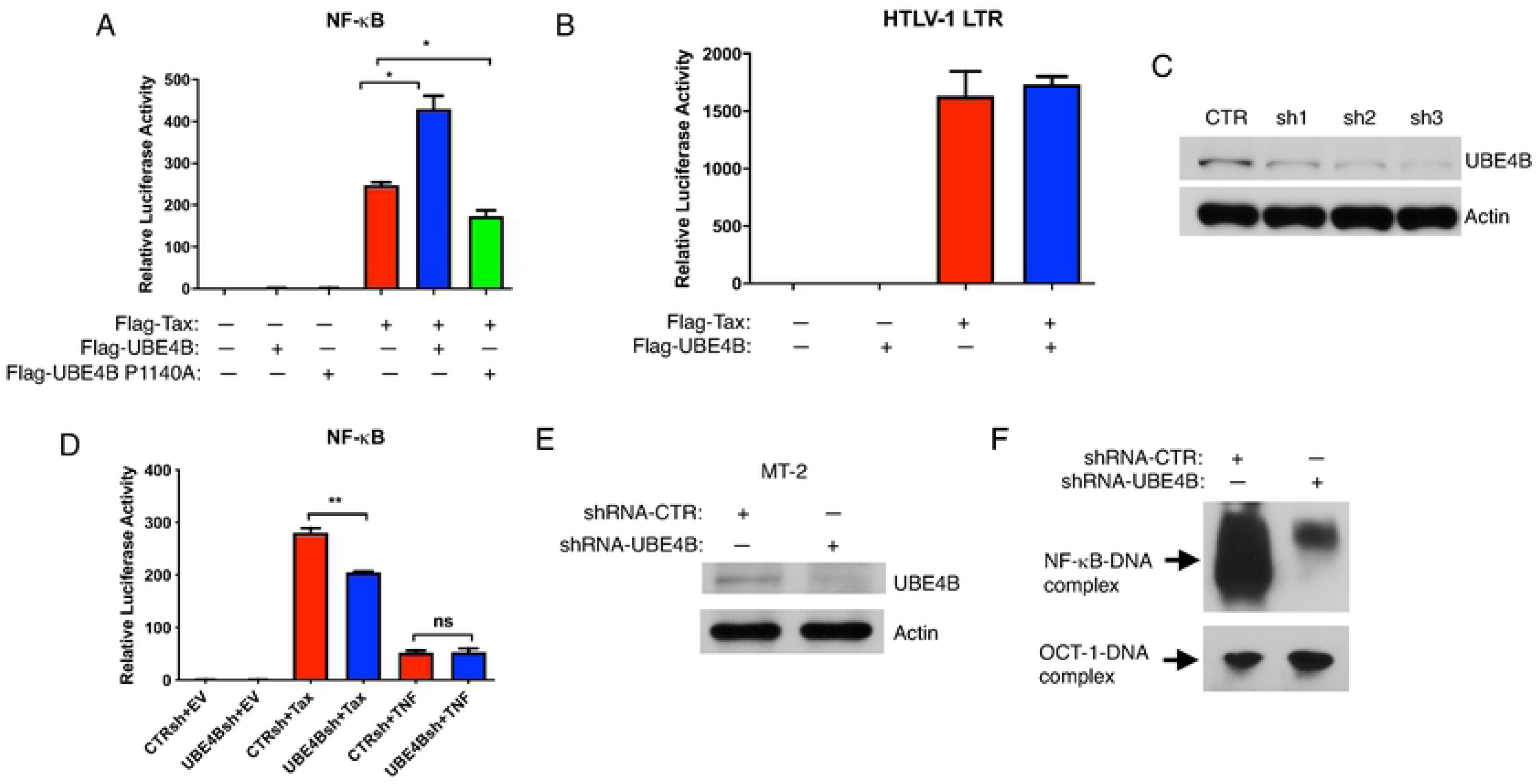
UBE4B enhances Tax-mediated activation of NF-κB. (A, B) NF-κB and HTLV-1 LTR luciferase assays in 293T cells transfected with NF-κB-TATA or HTLV-1 luciferase, pRL-tk, Flag-UBE4B, FLAG-UBE4B P1140A and Flag-Tax as indicated. (C) Immunoblotting was performed with the indicated antibodies using whole cell lysates from 293T cells expressing either control scrambled shRNA or UBE4B shRNAs 1-3. (D) NF-κB luciferase assays in 293T cells expressing either control shRNA or UBE4B shRNA were transfected with NF-κB-TATA luciferase, pRL-tk, Flag-Tax or stimulated with TNF for 8 h as indicated. (E) Immunoblotting was performed with the indicated antibodies using whole cell lysates from MT-2 cells expressing either control shRNA or UBE4B shRNA. (F) NF-κB and OCT-1 EMSA using nuclear extracts from MT-2 cells expressing either control shRNA or UBE4B shRNA.

To further examine the functional roles of the UBE4B-Tax interaction, UBE4B expression was suppressed with three independent shRNAs in 293T cells using recombinant shRNA expressing lentiviruses (Figure 3C). UBE4B shRNA3 was most effective at reducing UBE4B protein expression (Figure 3C), and was used for subsequent experiments. Tax-induced NF-κB activation was significantly decreased in 293T cells expressing UBE4B shRNA (Figure 3D), and the effects of UBE4B on NF-κB appeared to be specific for Tax since UBE4B depletion had no effect on tumor necrosis factor (TNF)-induced NF-κB activation (Figure 3D). To determine if UBE4B supported persistent NF-κB activation by Tax in HTLV-1 transformed cells, we knocked down UBE4B using lentiviral-expressed shRNA and performed an NF-κB DNA binding assay (EMSA) using nuclear extracts from MT-2 cells. Immunoblotting with lysates from MT-2 cells transduced with lentiviruses expressing control or UBE4B shRNAs, confirmed UBE4B knockdown (Figure 3E). As expected, robust NF-κB DNA binding was observed with MT-2 cells expressing control shRNA; however, shRNA-mediated knockdown of UBE4B in MT-2 cells significantly impaired NF-κB but not OCT-1 DNA binding (Figure 3F).

### Knockdown of UBE4B impairs Tax-induced expression of NF-κB target genes

Tax is required for the early steps of HTLV-1-mediated T-cell transformation, but may be dispensable at later times after development of ATLL. The HTLV-1 transformed cell lines MT-2, HUT-102 and C8166 have high levels of Tax expression and persistent NF-κB activation. Given that Tax expression is silenced in ∼60% of ATLL [12], we wondered if UBE4B supported NF-κB activation in Tax-negative ATLL cells. The ATLL cell line TL-OM1 is derived from an ATLL patient and lacks Tax expression. Although Tax is not expressed in these cells, persistent NF-κB activation is still maintained by genetic and/or epigenetic changes [42]. To confirm the effects of UBE4B on Tax-mediated NF-κB activation, the mRNA expression of NF-κB target genes was analyzed in Tax+ and Tax-ATLL cell lines by qRT-PCR after shRNA-mediated knockdown of UBE4B. UBE4B knockdown was confirmed by qRT-PCR in Tax+ MT-2, HUT-102, C8166 and Tax-TL-OM1 cell lines (Figure 4A). CD25 is the high affinity subunit of the IL-2 receptor and is critical for T-cell proliferation. UBE4B knockdown significantly impaired the expression of CD25 in Tax+ MT-2, HUT-102 and C8166 cell lines, but not in Tax-TL-OM1 cells (Figure 4B, C). Expression of IRF-4, a transcription factor critical for the survival of ATLL cell lines [43], was significantly decreased in MT-2, but not in HUT-102, C8166 or TL-OM1 cells upon UBE4B depletion (Figure 4B, C). Expression of cIAP-2, an anti-apoptotic NF-κB target gene, was significantly decreased in HUT-102, but not in MT-2, C8166 or TL-OM1 cells when UBE4B was knocked down (Figure 4B). It is unclear why UBE4B knockdown did not impact IRF-4 or cIAP-2 expression in all the Tax+ cell lines, but may reflect cell-type differences in regulation of these genes. Importantly, Tax expression was unaffected in MT-2, HUT-102 or C8166 cells upon UBE4B knockdown (Figure 4B).

**Figure 4.**
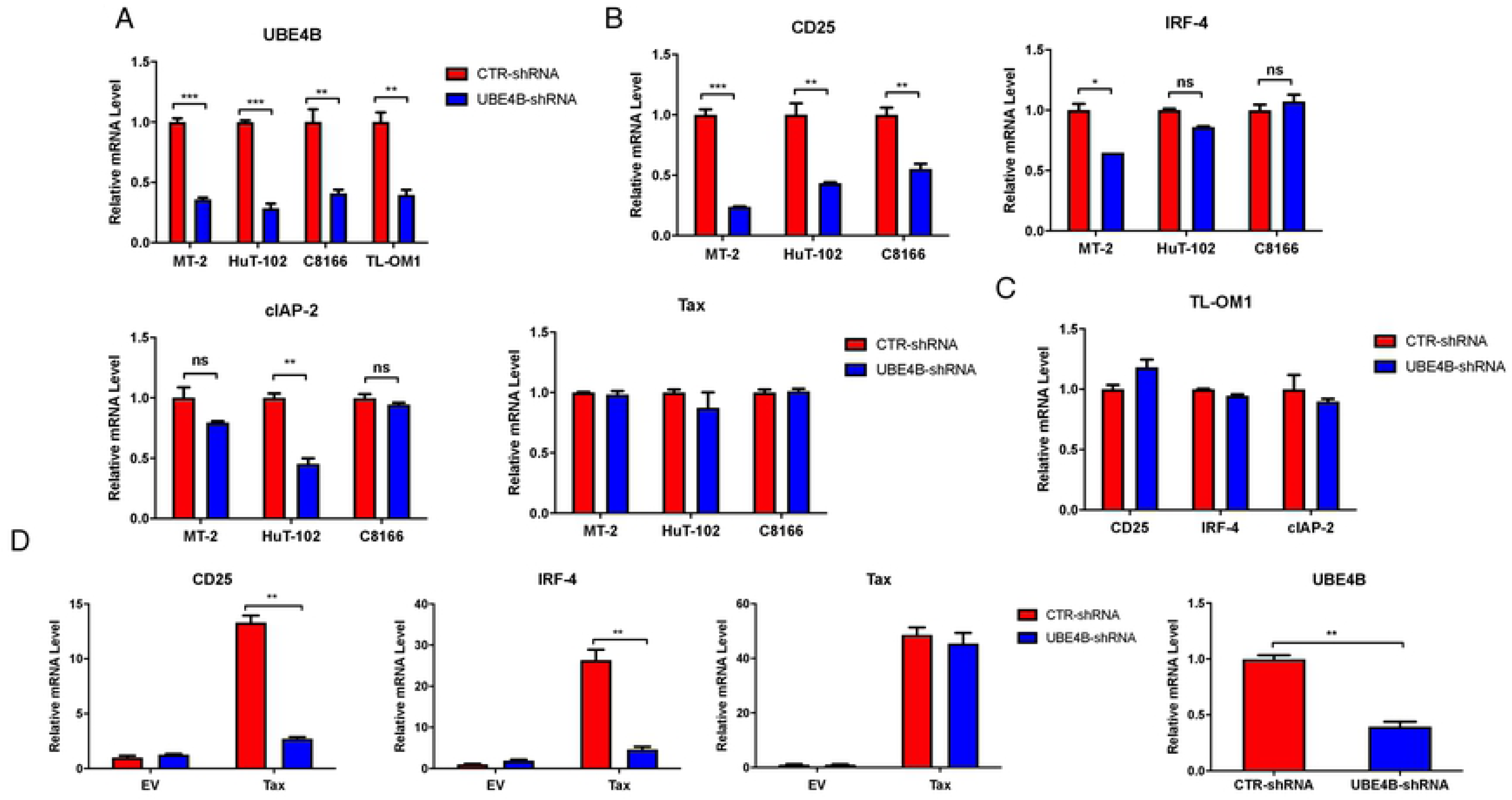
Knockdown of UBE4B impairs Tax-induced NF-κB target gene expression. (A-C) qRT-PCR of UBE4B, CD25, IRF-4, cIAP-2 and Tax mRNAs in MT-2, HUT-102, C8166 and TL-OM1 cells expressing control shRNA or UBE4B shRNA. (D) qRT-PCR of CD25, IRF-4, Tax and UBE4B mRNAs in Jurkat T cells expressing control shRNA or UBE4B shRNA and transduced with lentivirus expressing Tax.

To further confirm the effect of UBE4B on Tax-mediated NF-κB activation, UBE4B was knocked down in Jurkat T cells and Tax was introduced into the cells by lentiviral transduction. Cells were lysed two days later and mRNA was extracted for qRT-PCR experiments. The expression of both CD25 and IRF-4 were significantly reduced after UBE4B knockdown in Jurkat T cells expressing Tax (Figure 4D). However, Tax expression was comparable between the UBE4B knockdown and control samples (Figure 4D). UBE4B mRNA was significantly downregulated by the shRNA (Figure 4D). Taken together, UBE4B specifically mediates Tax-induced NF-κB activation and does not play a role in NF-κB activation in Tax-ATLL cells.

### UBE4B expression is not regulated by Tax or NF-κB

We considered the possibility that Tax may upregulate UBE4B expression to potentiate NF-κB signaling. To test this hypothesis, Jurkat Tax Tet-on T cells were treated with doxycycline (Dox) to induce Tax expression and after two days mRNA was extracted for qRT-PCR analysis. As expected, Tax upregulated the expression of CD25, however UBE4B expression was not increased by Tax (Figure S2A). Next, UBE4B expression was examined in a panel of HTLV-1 transformed cell lines that all exhibit persistent NF-κB activation. UBE4B mRNA (Figure S2B) and protein (Figure S2C) displayed variation in the panel of Tax+ (MT-2, HUT-102, C8166, MT-4), Tax- (TL-OM1, ED-40515(-) and ATL-43T) ATLL cell lines, as well as control Jurkat T cells and peripheral blood mononuclear cells (PBMCs) from a normal donor. However, the differences did not correlate with Tax expression (Fig. S2C). Therefore, UBE4B expression does not appear to be regulated by Tax or NF-κB.

### CRISPR/Cas9-mediated knockout of UBE4B impairs Tax-induced NF-κB activation

To further corroborate the UBE4B shRNA results on Tax-NF-κB activation, we next used CRISPR/Cas9 to generate UBE4B knockout (KO) 293T cells. Three different gRNAs targeting *Ube4b* exon 10 were expressed in pLentiCRISPRv2 for the production of recombinant lentiviruses expressing UBE4B gRNAs and Cas9. 293T cells were transduced with lentiviruses and the bulk population analyzed by immunoblotting for UBE4B knockout. Only one of the three gRNAs (gRNA3) resulted in efficient UBE4B knockout (data not shown), and therefore cells expressing either UBE4B gRNA3 or control gRNA were subjected to limiting dilution to isolate individual clones. Three individual KO clones (G12, H1 and F5) and two control clones (E2 and H10) were examined for UBE4B expression by immunoblotting and genomic DNA sequencing of *Ube4b* exon 10. UBE4B knockout clones G12 and H1 both harbored an insertion of an adenine in *Ube4b* exon 10 which would result in a frameshift mutation (Figure S3A). All three KO clones lacked expression of UBE4B as determined by immunoblotting (Figure S3B).

To examine Tax-induced NF-κB activation in UBE4B KO 293T cells, control and UBE4B KO clones were transfected with a Tax plasmid and subjected to NF-κB and HTLV-1 LTR luciferase assays. Tax activation of NF-κB was significantly reduced in all three UBE4B KO clones; however, Tax activation of the HTLV-1 LTR was either increased or unchanged in control and UBE4B KO clones (Figure 5A and B). Tax expression was comparable in each of the transfected clones (Figure 5A and B). We next examined phosphorylation of the NF-κB inhibitor IκBα using a phospho-specific antibody. Phosphorylation of IκBα by Tax was decreased in UBE4B KO cells suggesting that UBE4B functions at the level of or upstream of the IKK complex (Figure 5C).

**Figure 5.**
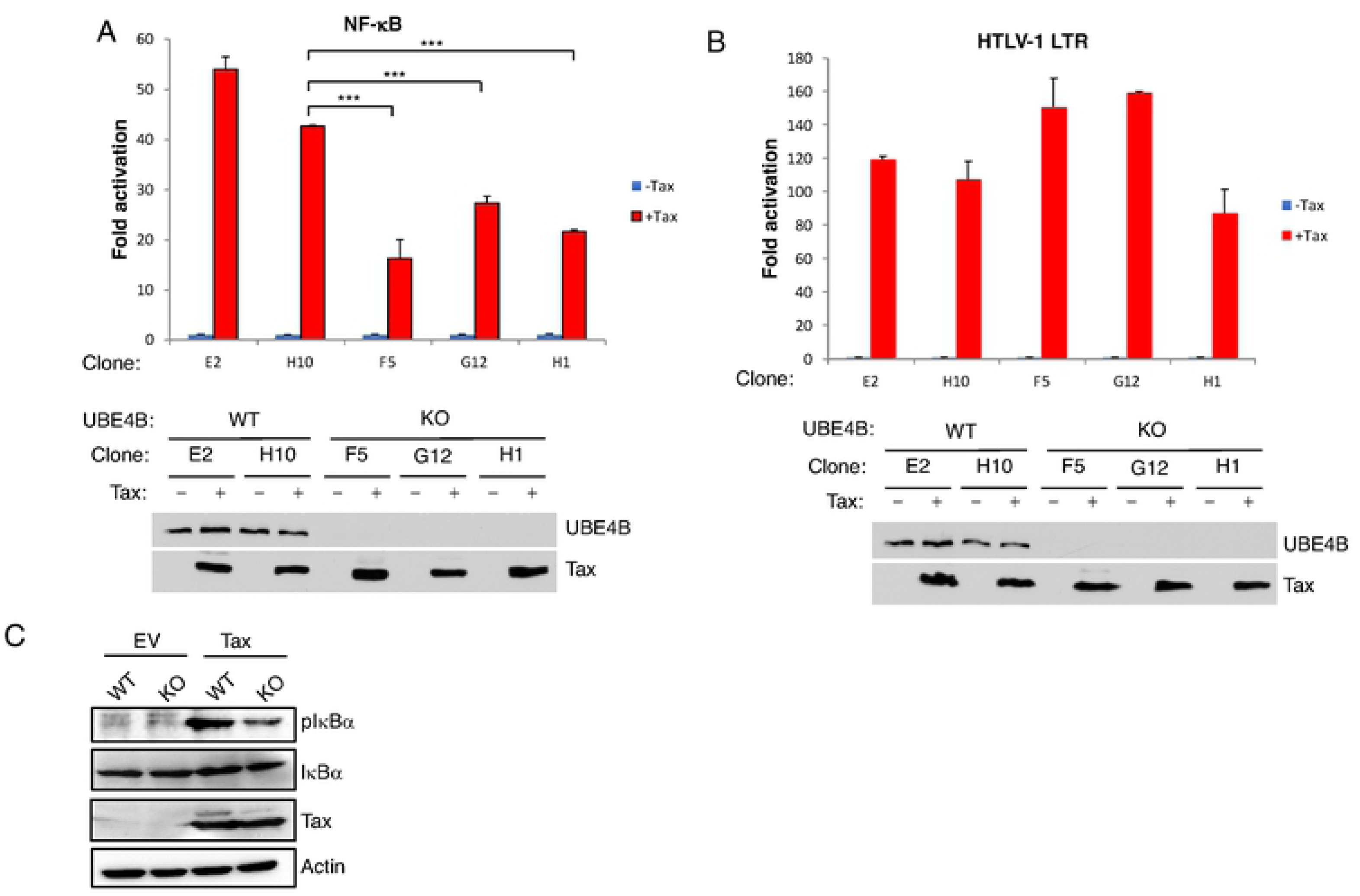
Tax activation of NF-κB is impaired in UBE4B KO cells. (A, B) NF-κB and HTLV-1 LTR luciferase assays in wild-type (E2, H10) and UBE4B KO (F5, G12, H1) 293T cells transfected with NF-κB-TATA or HTLV-1 LTR luciferase, pRL-tk, and Tax as indicated. Immunoblotting was performed with lysates from transfected cells. (C) Immunoblotting was performed with the indicated antibodies using lysates from wild-type and UBE4B KO (clone H1) 293T cells transfected with Tax.

### Knockdown of UBE4B promotes apoptosis in HTLV-1 transformed cell lines

Given that NF-κB is essential for HTLV-1-induced oncogenesis and cell survival, we next examined if UBE4B depletion with shRNA would trigger apoptosis of HTLV-1-transformed cells. Indeed, knockdown of UBE4B in MT-2 and HUT-102 cells, but not in control Jurkat cells, yielded cleaved forms of PARP and caspase 3 as detected by immunoblotting (Figure 6A), indicative of apoptotic cell death. We next examined the effect of UBE4B knockdown on the viability and proliferation of control Jurkat cells and the HTLV-1-transformed cell lines MT-2, HUT-102 and C8166 using CellTiter-Glo luminescent cell viability assay, which measures metabolically active cells by quantifying ATP levels. Jurkat, MT-2, HUT-102 and C8166 cells were infected with recombinant lentiviruses expressing either UBE4B or control shRNA. Metabolically active cells were quantified every 24 h for 5 consecutive days to measure cell proliferation. As expected, cells expressing control scrambled shRNA proliferated vigorously throughout the time course (Figure 6B-E). However, the proliferation of MT-2, HUT-102 and C8166, but not Jurkat cells, was significantly impaired upon UBE4B knockdown (Figure 6B-6E). Therefore, HTLV-1 transformed cell lines are dependent on UBE4B for proliferation and survival.

**Figure 6.**
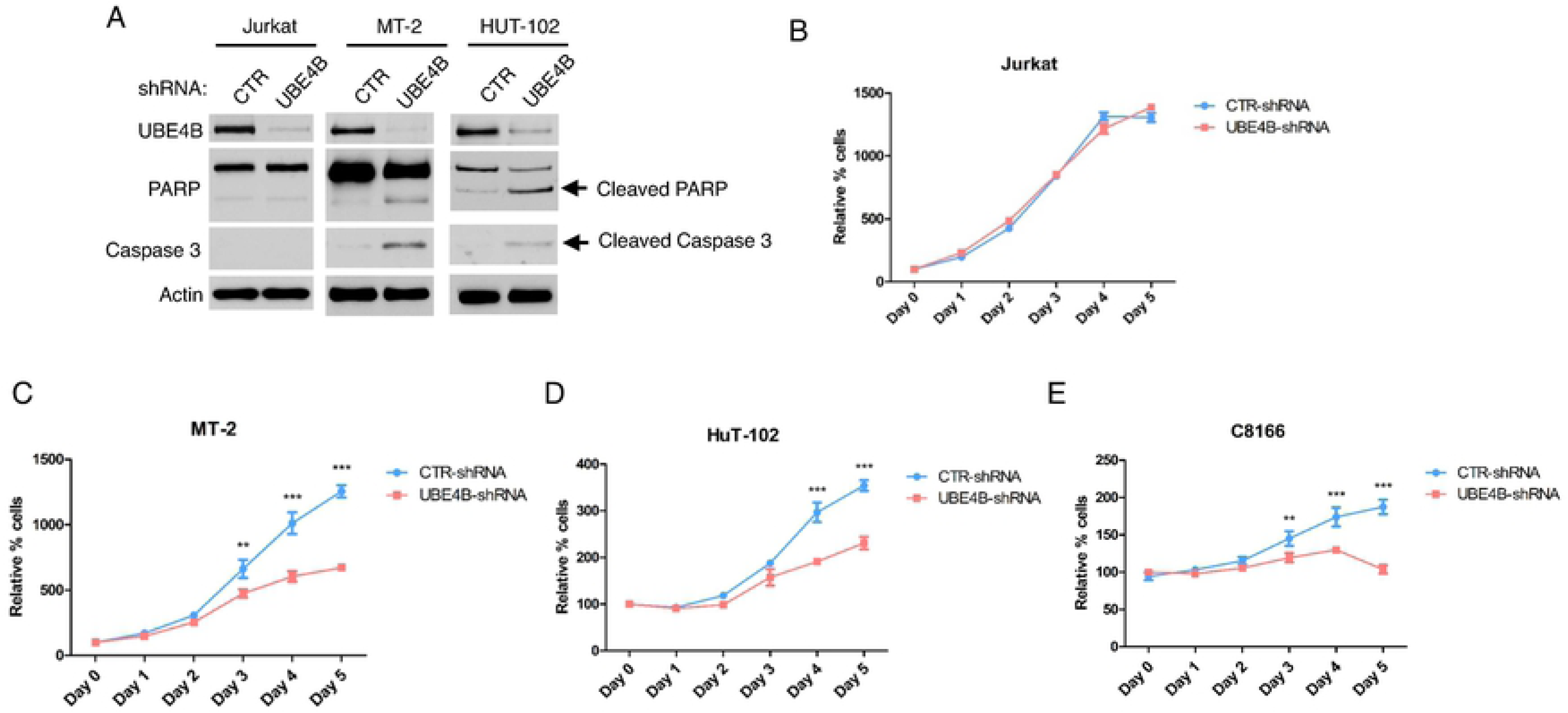
Knockdown of UBE4B promotes apoptotic cell death in HTLV-1 transformed cell lines. (A) Immunoblotting was performed with the indicated antibodies using lysates from Jurkat, MT-2 and HUT-102 cells expressing control shRNA or UBE4B shRNA. (B-E) Cell viability assay was performed at the indicated times with Jurkat, MT-2, HUT-102 and C8166 cells expressing control shRNA or UBE4B shRNA.

### UBE4B does not promote Rb and p53 degradation

The retinoblastoma (Rb) protein regulates a number of key cellular processes, including cell division, differentiation, senescence and apoptosis. It was reported that Tax can directly associate with and target Rb for proteasomal degradation [44]. UBE4B may potentially be recruited to Rb by Tax and function as an E3/E4 ligase to catalyze K48-linked polyubiquitination and degradation of Rb. To test this notion, UBE4B was knocked down with shRNA in Jurkat, MT-2, C8166 and HUT-102 cells, and lysates were subjected to immunoblotting to examine Rb expression. Consistent with the previous study, low levels of Rb protein were observed in HTLV-1 transformed cell lines MT-2, C8166 and HUT-102, compared to control Jurkat T cells (Figure S4). However, Rb protein expression was unaffected upon UBE4B knockdown in MT-2, C8166 and HUT-102 cells, indicating that UBE4B does not regulate Tax-induced Rb degradation (Figure S4).

UBE4B extends polyUb chains initiated by Hdm2 to induce the degradation of p53 [32]. To examine whether UBE4B regulated p53 in HTLV-1-transformed cell lines, p53 protein expression was monitored by immunoblotting after UBE4B knockdown. However, the expression of p53 protein was unchanged in MT-2, C8166 and HUT-102 cells after UBE4B knockdown (Figure S4). Therefore, UBE4B does not appear to regulate p53 stability in HTLV-1 transformed cell lines, therefore p53 ubiquitination by UBE4B may be cell-type specific or Tax uses distinct mechanisms to inhibit p53.

### UBE4B is required for Tax polyubiquitination

Since UBE4B is an E3/E4 ubiquitin ligase and it potentiates Tax-mediated NF-κB activation, we next examined whether overexpression of UBE4B promotes Tax ubiquitination. 6xHis-Tax was transfected into cells, with and without Flag-UBE4B, and lysates were used to purify Tax with Ni-NTA agarose. Tax polyUb chains were assessed by immunoblotting for total ubiquitin (Ub) and K63-linked Ub. Overexpressed UBE4B enhanced Tax total polyUb and K63-linked polyUb chains (Figure 7A). The catalytic activity of UBE4B was required since the UBE4B P1140A mutant was impaired in enhancing Tax polyUb chains (Figure 7B). We next performed loss of function studies using 293T cells expressing Flag-Tax, and either control shRNA or UBE4B shRNA. Both total and K63-linked polyubiquitination of Tax were impaired when UBE4B expression was suppressed (Figure 7C). Thus, UBE4B positively regulates Tax polyubiquitination, including K63-linked polyUb chains.

**Figure 7.**
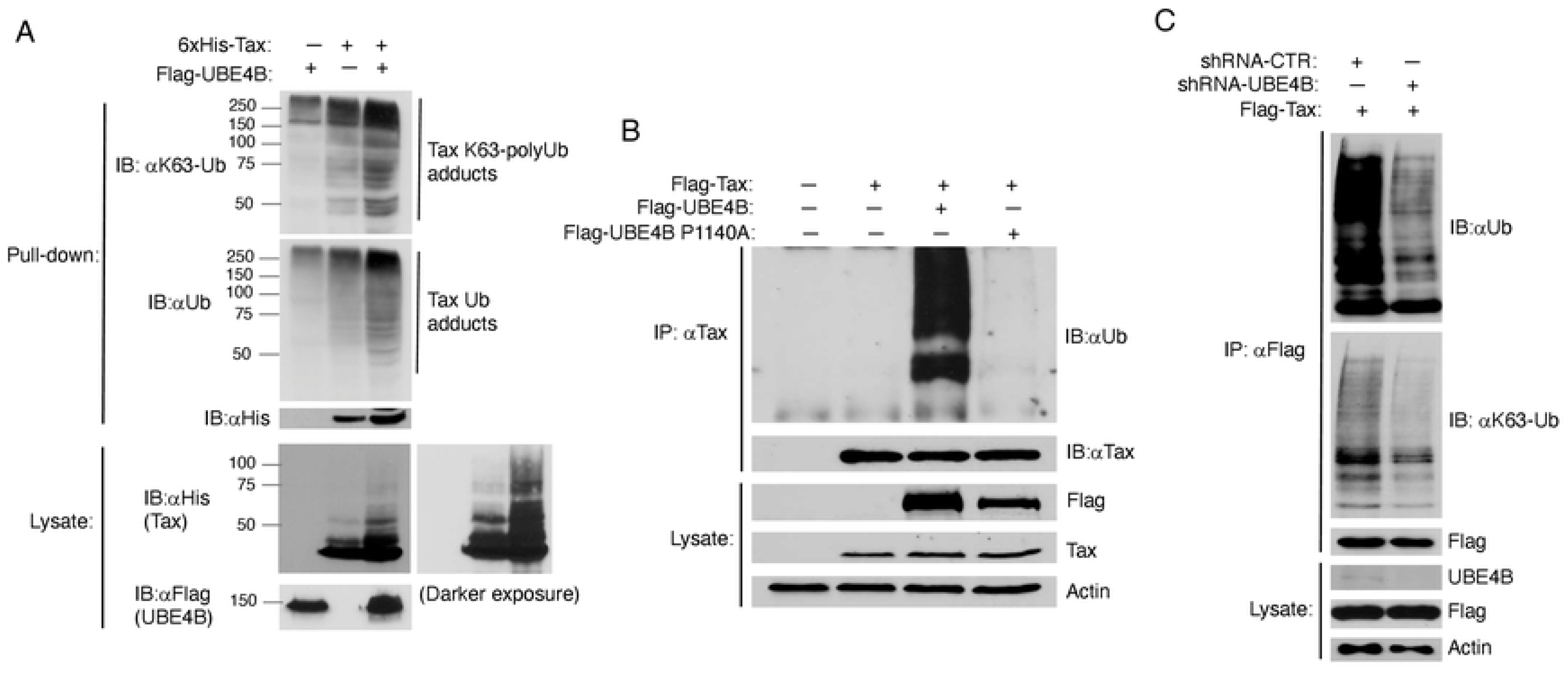
Tax promotes the polyubiquitination of UBE4B. (A) Ubiquitination assay was performed with purified 6xHis-Tax from lysates of 293T cells transfected with 6xHis-Tax and Flag-UBE4B as indicated. Immunoblotting was performed with the indicated antibodies. (B) Ubiquitination assay was performed with Tax immunoprecipitates from lysates of 293T cells transfected with Flag-Tax, Flag-UBE4B and Flag-UBE4B P1140A. (C) Ubiquitination assay was performed with Flag-Tax immunoprecipitates from lysates of 293T cells expressing control shRNA or UBE4B shRNA and transfected with Flag-Tax.

We further examined Tax polyUb in UBE4B KO 293T cells. Tax was expressed in control and UBE4B KO cells, and protein lysates were used for Tax ubiquitination assays. In agreement with our earlier shRNA knockdown experiments, Tax total and K63-linked polyUb chains were decreased in UBE4B KO cells (Figure 8A, B). We also found that Tax K48-linked polyUb chains were sharply reduced in UBE4B KO cells (Figure 8B). However, Tax stability, as determined by a cycloheximide chase assay, was similar in WT and UBE4B KO cells (Figure S5). Therefore, UBE4B does not appear to promote Tax degradation, but is critical for both K48- and K63-linked Tax polyUb chains.

**Figure 8.**
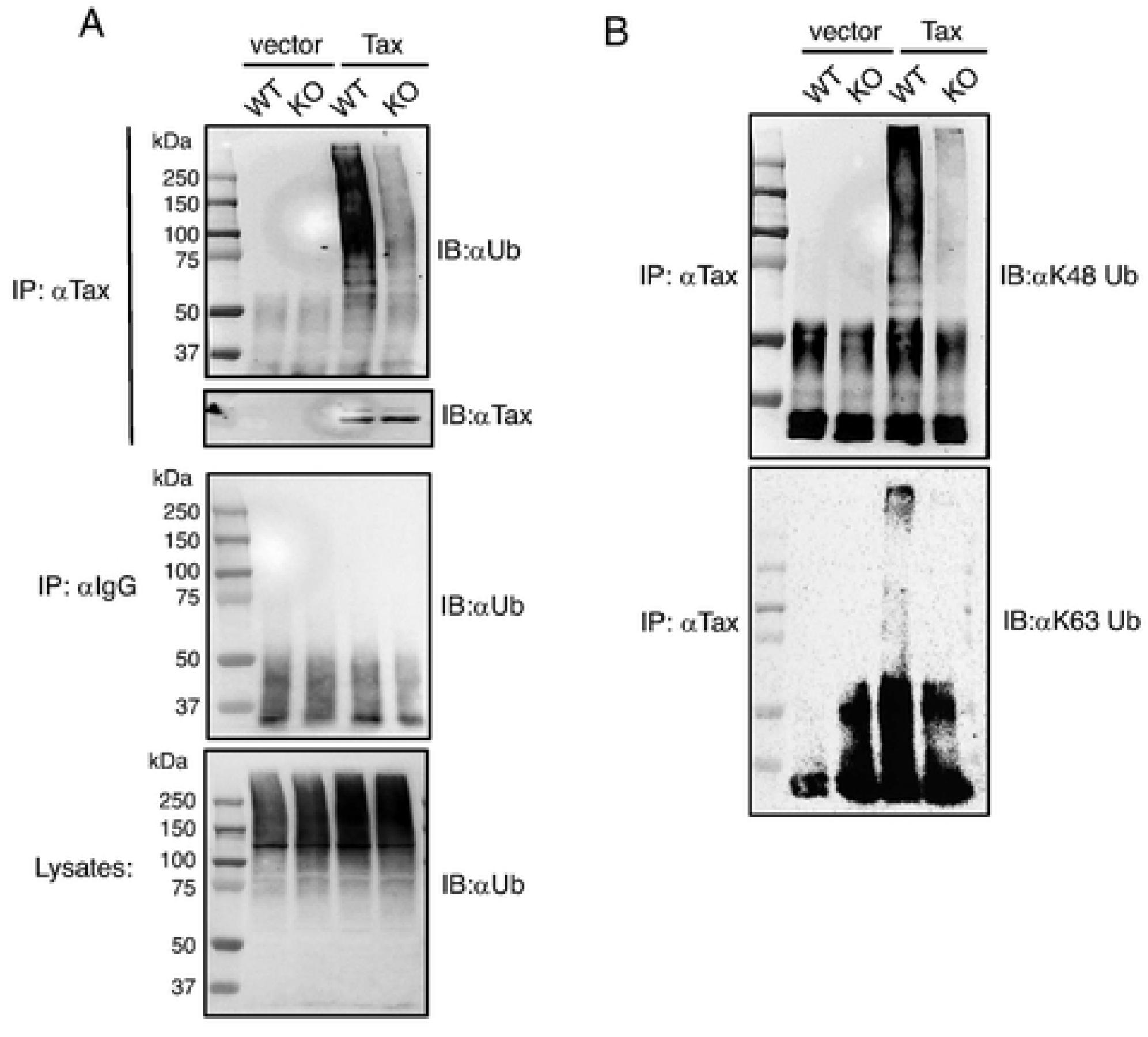
Tax polyubiquitination is impaired in UBE4B KO cells. (A, B) Ubiquitination assays were performed with Tax immunoprecipitates from lysates of wild-type and UBE4B KO (Clone H1) 293T cells transfected with Tax as indicated.

Since UBE4B and Tax can interact, and UBE4B supports Tax polyubiquitination, we next determined if UBE4B could directly ubiquitinate Tax. To this end, we performed an *in vitro* ubiquitination assay with recombinant E1, E2 (UbcH5c), Ub and UBE4B-His6 proteins as well as immunopurified Tax from 293T cell lysates. After the *in vitro* Ub assay, Tax was subjected to a second IP followed by immunoblotting with anti-Ub. Tax polyUb chains were observed only in the presence of E1, E2, UBE4B, Ub and Tax (Figure 9). Therefore, we conclude that UBE4B can directly conjugate Tax with polyUb chains.

**Figure 9.**
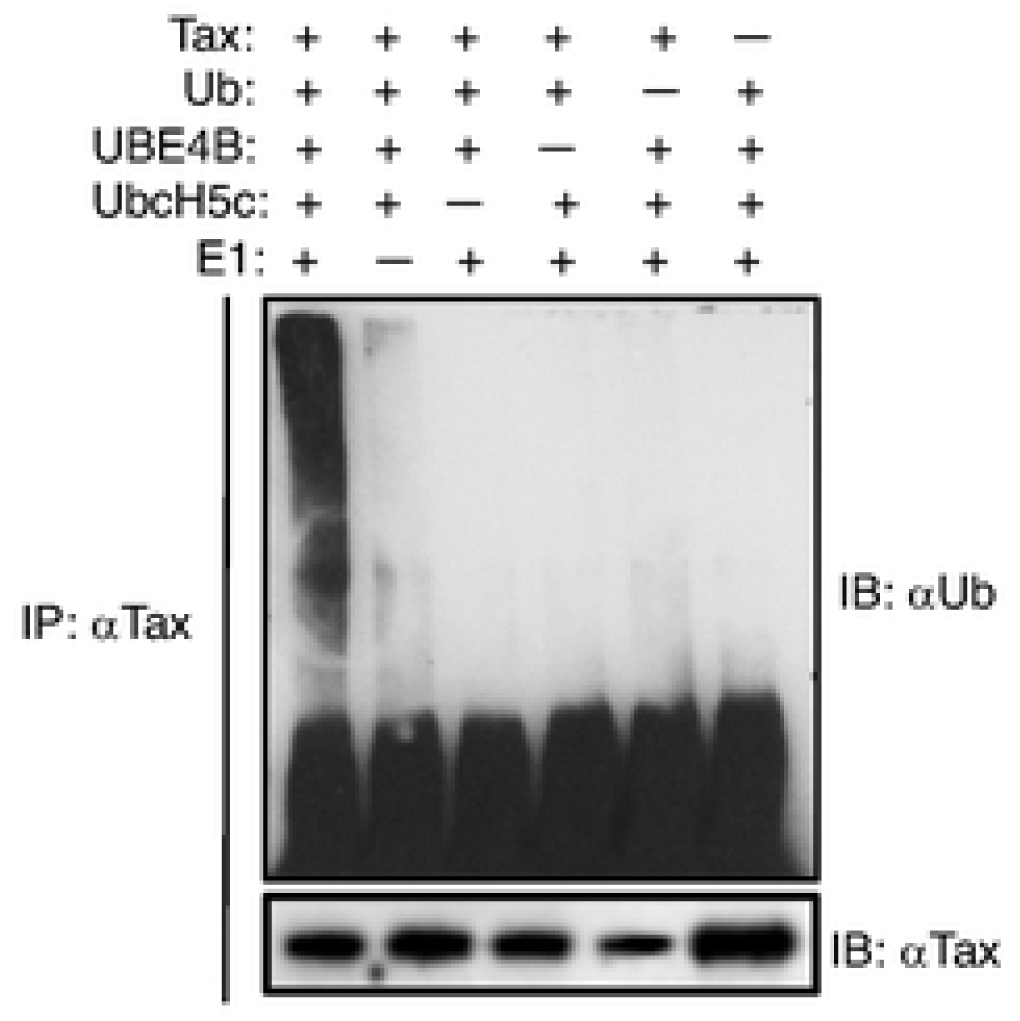
UBE4B directly ubiquitinates Tax. *In vitro* ubiquitination assay was performed with the indicated recombinant proteins and immunoprecipitated Tax from lysates of transfected 293T cells. Following the Ub assay, Tax was immunoprecipitated from the reaction mixtures and immunoblotted with the indicated antibodies.

## Discussion

Our study has identified the E3/E4 ubiquitin conjugation factor UBE4B as a novel interacting protein of HTLV-1 Tax. We have confirmed using multiple experimental approaches that UBE4B interacts with and colocalizes with Tax in HTLV-1 transformed T cells. Knockdown or knockout of UBE4B impairs Tax-induced NF-κB activation, as well as NF-κB signaling and cell survival in Tax+ HTLV-1 transformed cells. Furthermore, overexpression of UBE4B enhances Tax polyubiquitination, whereas loss of UBE4B impairs Tax K48- and K63-linked polyUb chains. Finally, we have demonstrated that UBE4B directly conjugates Tax with polyUb chains. Collectively, these results support the model depicted in Figure 10 where UBE4B interacts with and ubiquitinates Tax to activate downstream NF-κB signaling.

**Figure 10.**
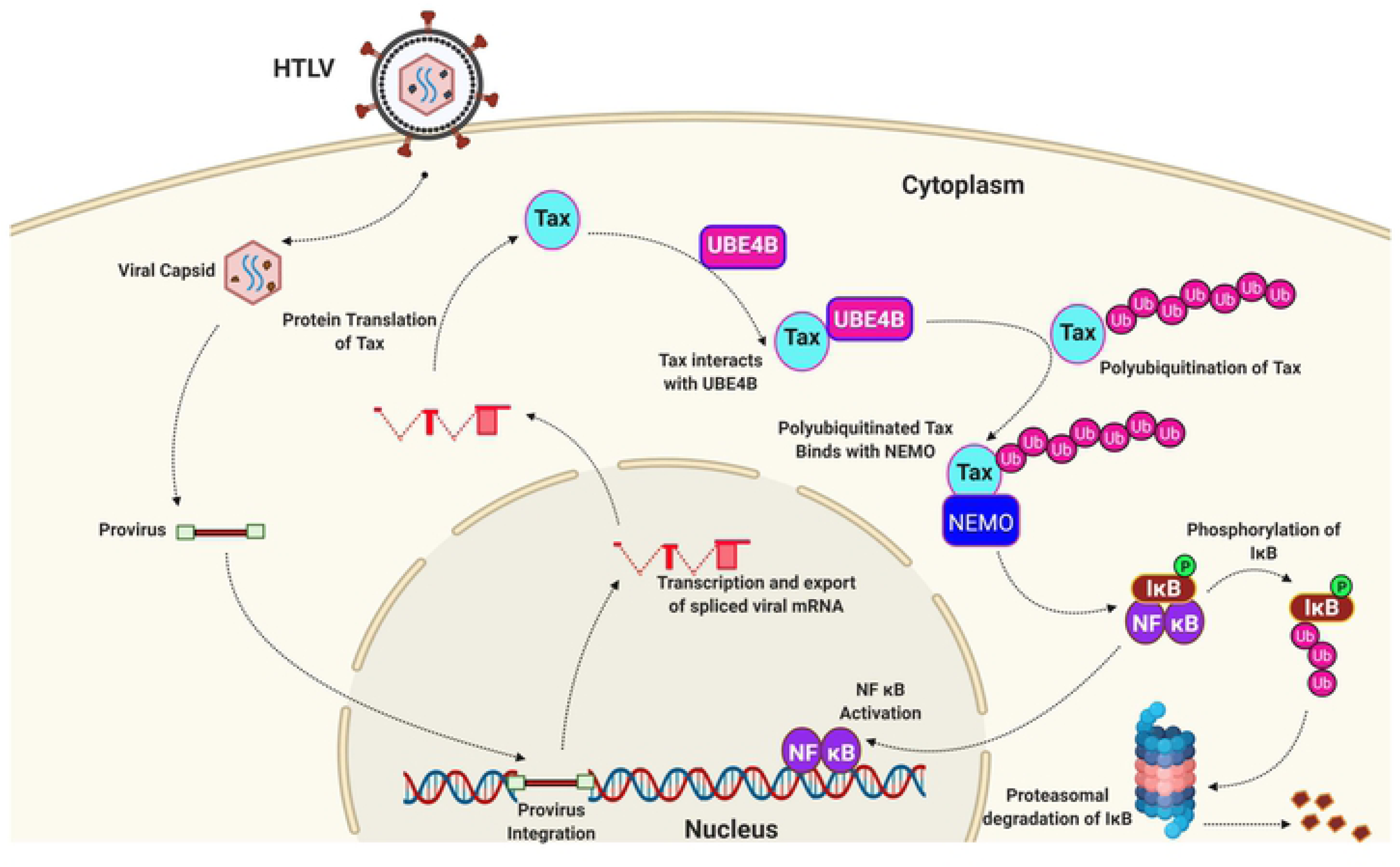
Model of UBE4B regulation of Tax polyubiquitination and NF-κB activation. Tax interacts with UBE4B, and UBE4B promotes Tax polyUb for downstream IKK and NF-κB activation.

Our confocal microscopy and PLA experiments suggest that UBE4B and Tax mainly co-localize in the cytoplasm, however nuclear UBE4B-Tax complexes were also detected suggesting that Tax-UBE4B complexes may shuttle between the cytoplasm and nucleus. Tax can localize within lipid rafts near the cis-Golgi where it directs relocalization and activation of the IKK complex [15, 45]. Since Tax K63-linked polyubiquitination is necessary to relocalize IKK to the cis-Golgi, UBE4B may be important for Tax to recruit IKK to the cis-Golgi. Tax-induced IκBα phosphorylation was impaired in UBE4B KO cells (Figure 5C), consistent with UBE4B positioned upstream of the IKK complex together with Tax. Furthermore, UBE4B appears to selectively mediate Tax-induced NF-κB activation, but not TNF-triggered NF-κB suggesting a specific role of UBE4B in the regulation of Tax.

Knockdown of UBE4B triggered apoptotic cell death and diminished the proliferation of HTLV-1 transformed cell lines (Figure 6). Since UBE4B promotes Hdm2-mediated p53 proteasomal degradation in the nervous system [32], and Tax induces Hdm2-mediated Rb ubiquitination and subsequent degradation [44] we examined a potential regulation of p53 and Rb by UBE4B in HTLV-1 transformed T cells. However, knockdown of UBE4B had no influence on p53 and Rb protein expression in HTLV-1 transformed cell lines (Figure S4). Therefore, it appears that the effects of UBE4B on the proliferation and survival of HTLV-1 transformed cells are mediated predominantly through NF-κB signaling. However, we cannot rule out additional effects of UBE4B on the cell cycle, ER stress/ERAD, or other pathways that impinge on cell proliferation and survival.

UBE4B supports both K63- and K48-linked polyubiquitination of Tax (Figures 7 and 8). Although it has not been demonstrated that UBE4B can catalyze K63-linked polyUb chains, this does not rule out the possibility that UBE4B conjugates Tax with K63-linked polyUb chains. The specificity of the polyUb linkage catalyzed by a U-box ligase is determined by the E2 ubiquitin-conjugating enzyme, rather than the E3 enzyme. Furthermore, UBE4B can utilize UbcH5c as an E2 enzyme, which has the potential to catalyze K63-linked polyUb chains [46]. Our *in vitro* ubiquitination assay (Figure 9) supports the notion that UBE4B directly ubiquitinates Tax; however, we cannot distinguish if the E3 and/or E4 activity of UBE4B contributes to Tax polyubiquitination. Immunopurified Tax from cell lysates likely has post-translational modifications including mono- and polyUb which could potentially be extended and fine-tuned by the E4 activity of UBE4B. Therefore, we envision two potential mechanisms of UBE4B regulation of Tax polyUb: 1) UBE4B functions as an E3 ligase for Tax and conjugates Tax with both K48- and K63-linked polyUb chains, or 2) UBE4B functions as an E4 enzyme and extends K48- and/or K63-linked polyUb chains primed by other E3 enzymes. Tax itself has been proposed to have E3 ligase activity [47], however we did not observe Tax polyUb chains in the absence of UBE4B in an *in vitro* Ub assay (Figure 9).

It is curious that UBE4B promotes K48-linked polyubiquitination of Tax but does not appear to trigger its degradation. Based on these findings, one possibility is that UBE4B may mediate branched K48-K63 polyUb chains on Tax. In this regard, it has been reported that the E3 ligase TRAF6 is conjugated with branched K48-K63 polyUb chains by the E3 ubiquitin ligase HUWE1, and these branched chains enhance TRAF6-mediated NF-κB activation by inhibiting the disassembly of K63-linked polyUb chains by deubiquitinating enzymes such as CYLD [48]. Similarly, UBE4B may potentially catalyze branched K48- K63 polyUb chains on Tax to stabilize and protect the K63-linked polyUb chains from negative regulators, and therefore potentiate persistent NF-κB signaling. There is precedence for E4 enzymes in synthesizing branched Ub chains as yeast Ufd2p can catalyze K29-K48 branched polyUb chains to promote degradation of substrates [49]. Additional experiments are needed to further investigate if Tax is conjugated with branched K48-K63 polyUb chains, and if UBE4B is implicated in this process.

Overall, we have identified the E3/E4 conjugation factor UBE4B as a Tax binding partner that promotes Tax polyubiquitination and NF-κB activation. Since Tax K63-linked polyubiquitination is required for constitutive NF-κB activation and subsequent immortalization and transformation of CD4+ T cells, UBE4B may represent a novel therapeutic target for early stage HTLV-1-induced leukemogenesis and also Tax+ ATLL tumors (∼40% of all ATLL tumors express Tax).

## Materials and Methods

### Ethics statement

Blood from healthy donors was purchased from Biological Specialty Corporation (Colmar, PA). PBMCs were isolated as described previously [50].

### Reagents, plasmids and antibodies

Human embryonic kidney cells (HEK 293T) were purchased from ATCC. Cell lines Jurkat, Jurkat Tax Tet-on, C8166, MT-2, MT-4, HUT-102, TL-OM1, ED40515(-) and ATL43T were described previously [50, 51]. HEK 293T cells were cultured in Dulbecco’s modified Eagle’s medium (DMEM); Jurkat, Jurkat Tax Tet-on, C8166, MT-2, MT-4, HUT-102, TL-OM1, ED40515(-) and ATL43T cells were cultured in RPMI medium. Medium was supplemented with fetal bovine serum (10%) and penicillin-streptomycin (1%). Expression vectors encoding Flag-Tax, pCMV4-Tax, Tax M22, Tax M47, HTLV-1 LTR-Luc, NF-κB-TATA Luc, pRL-tk, pDUET-Tax, psPAX2 and VSV-G were described previously [17, 25, 52]. pDEST51-UBE4B-Flag was a gift from Dr. Sarah Spinette [53]. Site-directed mutagenesis of UBE4B (P1140A) was generated by PCR using Platinum Pfx DNA polymerase (Thermo Fisher Scientific). The monoclonal anti-Tax antibody was prepared from a Tax hybridoma (168B17-46-34) received from the AIDS Research and Reference Program, NIAID, National Institutes of Health. Anti-Tax antibody (1A3) was purchased from Santa Cruz Biotechnology. Alexa Fluor 594-conjugated donkey anti-mouse IgG and Alexa Fluor 488-conjugated donkey anti-rabbit IgG were purchased from Thermo Fisher Scientific. The monoclonal Flag M2 and hemagglutinin (HA; 12CA5) antibodies were purchased from Millipore-Sigma. UBE4B antibodies were purchased from Bethyl Laboratories and Santa Cruz Biotechnology. The NEMO, LDH, Rb, p53 and Caspase3 antibodies were from Santa Cruz Biotechnology. PARP, pIκBα, IκBα, Vinculin, β-Actin, K48- and K63 linkage-specific Ub antibodies were from Cell Signaling Technology. Ubiquitin antibody was purchased from Enzo Life Sciences. DAPI (4’, 6-diamidino-2-phenylindole) was purchased from EMD Biosciences. TNF was purchased from R&D Systems. Cycloheximide solution was purchased from Millipore-Sigma.

### Transfections and luciferase reporter assays

293T cells were transiently transfected with GenJet^TM^ In Vitro DNA Transfection Reagent (SignaGen Laboratories). Luciferase reporter assays were performed 24 h after DNA transfection, unless otherwise indicated, using the Dual-Glo luciferase assay system (Promega). Firefly luciferase values were normalized based on the *Renilla* luciferase internal control values. Luciferase values are presented as “fold induction” compared to the control transfected with empty vector.

### Immunoblotting, co-immunoprecipitation and ubiquitination assays

Whole cell lysates were generated by lysing cells in RIPA buffer (50 mM Tris-Cl [pH 7.4], 150 mM NaCl, 1% NP-40, 0.25% sodium deoxycholate, 1 mM phenylmethylsulfonyl fluoride [PMSF], 1× Roche complete mini-protease inhibitor cocktail) on ice, followed by centrifugation. Cell lysates were resolved by SDS-PAGE, transferred to nitrocellulose membranes, and subjected to immunoblotting with the indicated primary antibodies and HRP-conjugated secondary antibodies (GE Healthcare Life Sciences). Immunoreactive bands were visualized by Western Lightning enhanced chemiluminescence (PerkinElmer). For co-IPs, lysates were diluted 1:1 in RIPA buffer and precleared with protein A agarose beads (Millipore-Sigma) for 60 min at 4°C. Pre-cleared lysates were further incubated at 4°C overnight with the indicated antibodies (1 to 3 μl) and protein A agarose or protein G Dynabeads (Thermo Fisher Scientific). Immunoprecipitates were washed three times with RIPA buffer (LSB) to elute bound proteins. An additional wash with RIPA buffer supplemented with 1 M urea was performed for ubiquitination assays. For Tax ubiquitination assays performed with 6xHis-Tax, cells were lysed in buffer B (100 mM NaH_2_PO_4_, 10 mM Tris, and 8 M urea [pH 8.0]) and His-tagged Tax proteins were precipitated with Ni-nitrilotriacetic acid (NTA) agarose (Qiagen). After washing in buffer C (100 mM NaH_2_PO_4_, 10 mM Tris, and 8 M urea [pH 6.3]), His-tagged proteins were eluted in buffer E (100 mM NaH_2_PO_4_, 10 mM Tris, and 8 M urea [pH 4.5]) and subjected to SDS-PAGE and immunoblotting.

### EMSA DNA binding assay

Nonradioactive EMSA was performed as described previously [50] using LightShift Chemiluminescent EMSA Kit (Thermo Scientific) according to the manufacturer’s instructions.

### Confocal microscopy

MT-2 and C8166 cells were cultured for 4 h on glass coverslips coated with poly-L-lysine in 6-well plates. Cells were fixed with 4% paraformaldehyde for 30 min and permeabilized with 0.5% Triton X-100 for 5 min. The fixed cells were then incubated with 5% BSA for 1 h followed by staining with mouse anti-Tax or rabbit anti-UBE4B antibodies overnight at 4°C. Coverslips were incubated with Alexa Fluor 594-conjugated donkey anti-mouse IgG, Alexa Fluor 488-conjugated donkey anti-rabbit IgG (Thermo Fisher Scientific), and DAPI to stain nuclei. Images were acquired with a C2+ confocal microscope system (Nikon), and processed using NIS Elements software.

### Proximity ligation assay (PLA)

PLA was performed with the DuoLink^TM^ In Situ Red Starter Kit Mouse/Rabbit (Millipore-Sigma) as recommended by the manufacturer. MT-2 and C8166 cells were grown on glass coverslips, fixed, permeabilized and incubated with primary antibodies: anti-UBE4B (Rabbit) or anti-Tax (Mouse). Slides were incubated with Duolink PLA probes, ligated, amplified and washed. Images were acquired with a C2+ confocal microscope system (Nikon). PLA signals were quantified using ImageJ software.

### *In vitro* ubiquitination assay

Recombinant E1 ubiquitin activating enzyme, E2 (UbcH5c), Ub, and UBE4B-His6 were purchased from BostonBiochem/R&D Systems. Tax was transfected into 293T cells and lysates immunoprecipitated with anti-Tax. Ub assay was performed on eluted immunoprecipitated Tax after addition of E1, UbcH5c and UBE4B-His6, and ubiquitin conjugation reaction buffer for 2 h at 37°C. Tax was immunoprecipitated again with anti-Tax followed by immunoblotting with anti-Ub.

### Quantitative real-time PCR (qRT-PCR)

RNA was isolated using the RNeasy minikit (Qiagen). RNA was converted to cDNA using the First Strand cDNA synthesis kit for reverse transcription (avian myeloblastosis virus [AMV]; Millipore-Sigma). Quantitative real-time PCR (qRT-PCR) was performed with an Applied Biosystems 7500 Real-Time PCR system using KiCqStart®SYBR®Green qPCR ReadyMix™ (Millipore-Sigma). Gene expression was normalized to the internal control 18S rRNA. Primer sequences for qRT-PCR are provided in Table S1.

### CHX chase assay

Cycloheximide (CHX) chase assays were performed as described previously [25]. Cells were treated with CHX (10 μg/ml) for various times 2 days after transfection. Cells were lysed in RIPA buffer, and immunoblotting was conducted with anti-Tax.

### Knockdown with lentiviral shRNAs

Three lentiviral Mission short hairpin RNA (shRNA) clones targeting UBE4B were obtained from Millipore-Sigma. HEK293T cells were transfected with the lentiviral shRNA-targeting vectors, psPAX2 packaging plasmid (Addgene) and vesicular stomatitis virus glycoprotein (VSV-G). After 48 h, the supernatants were collected and centrifuged at 25,000 rpm using a Beckman SW28 centrifuge rotor for 2 h at 4°C. The supernatants were removed, and the pellets were re-suspended in ice-cold phosphate-buffered saline (PBS). Viral stocks were used to infect cell lines.

### CRISPR/Cas9 knockout

CRISPR/Cas9-mediated knockout of UBE4B was performed as previously described [54]. Briefly, gRNAs specific for UBE4B exon 10 were designed through the E-CRISP web server (http://www.e-crisp.org/E-CRISP/) and expressed by pLentiCRISPRv2-puro (a gift from Feng Zhang; Addgene). Guide sequence function was analyzed using the Surveyor Mutation Detection Kit (Integrated DNA Technologies) according to the manufacturer’s instructions. The genomic DNA fragments including gRNA target sequences were PCR-amplified and analyzed by agarose gel electrophoresis. Individual clones were isolated by limiting dilution and genomic DNA purified, *Ube4b* exon 10 amplified by PCR and subjected to Sanger DNA sequencing. Primer sequences for gRNAs and Surveyor primers are provided in Table S1.

### Cell viability and proliferation assays

Cell viability was determined using the CellTiter-Glo luminescent cell viability assay (Promega), which quantitates ATP as a measure of metabolically active cells. A total of 50 μl of suspended cells and 50 μl of CellTiter-Glo solution were mixed and incubated at room temperature for 10 min, and the luminescence was quantified with a GloMax96 microplate luminometer (Promega).

### Yeast Two-Hybrid Analysis

Yeast two-hybrid screening was performed by Hybrigenics Services, S.A.S., Paris, France (http://www.hybrigenics-services.com). The coding sequence for the full length Tax protein was PCR-amplified and cloned into pB27 as a C-terminal fusion to LexA (N-LexA-Tax-C). The construct was checked by sequencing the entire insert and used as a bait to screen a random-primed human leukocyte and activated mononuclear cells cDNA library constructed into pP6. pB27 and pP6 derive from the original pBTM116 [55] and pGADGH [56] plasmids, respectively. 46.7 million clones were screened using a mating approach with YHGX13 (Y187 ade2-101::loxP-kanMX-loxP, matα) and L40ΔGal4 (mata) yeast strains as previously described [57]. 282 His+ colonies were selected on a medium lacking tryptophan, leucine and histidine. The prey fragments of the positive clones were amplified by PCR and sequenced at their 5’ and 3’ junctions. The resulting sequences were used to identify the corresponding interacting proteins in the GenBank database (NCBI) using a fully automated procedure. A confidence score (PBS, for Predicted Biological Score) was attributed to each interaction as previously described [58].

### Statistical analysis

Data are expressed as mean fold increase ± standard deviation relative to the control from a representative experiment performed 3 times in triplicate. Statistical analysis was performed in GraphPad Prism 8. * *P* value of <0.05, ** *P* value of <0.01, *** *P* value of <0.001 as determined by Student’s *t* test.

## Acknowledgements

We thank Dr. Sarah Spinette for the Flag-UBE4B plasmid and Dr. Masao Matsuoka for ATLL cell lines.

## Supplementary Figure Legends

**Figure S1.**
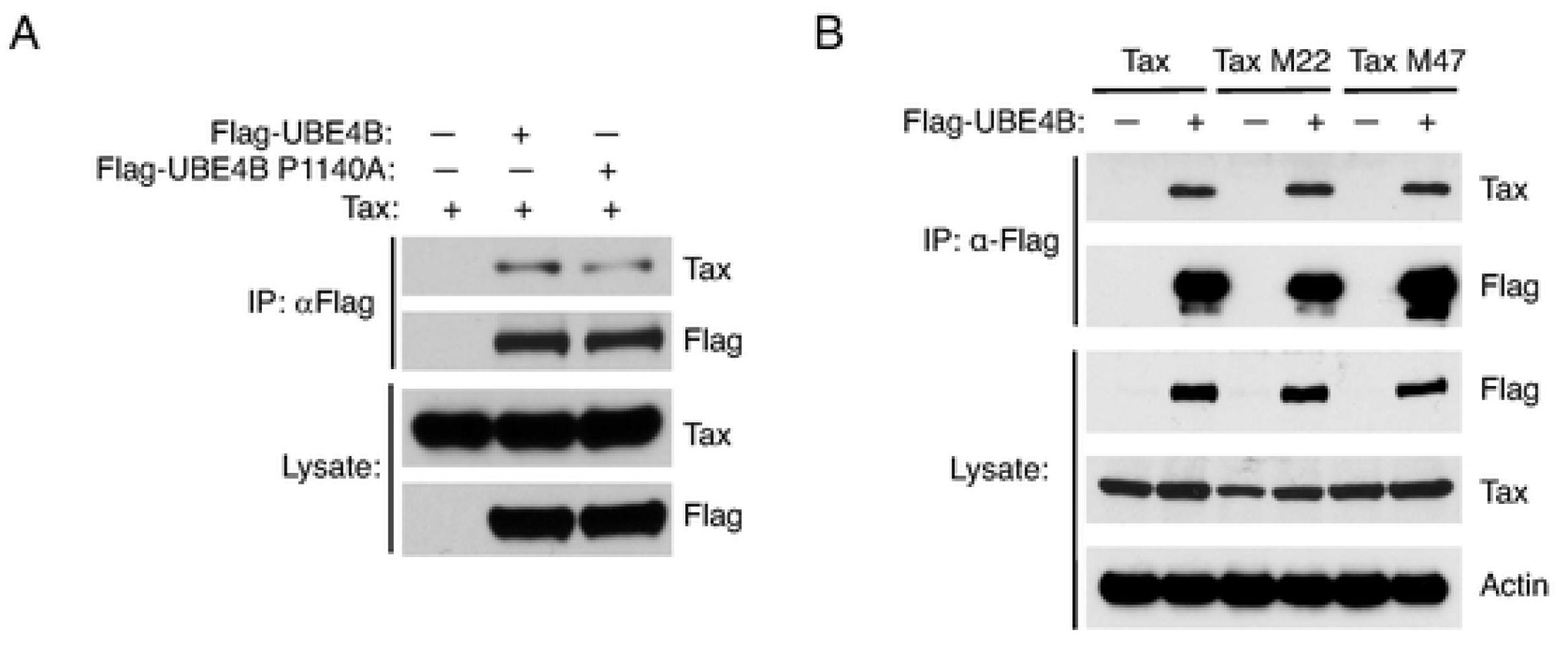
Interaction of Tax and UBE4B mutants. (A, B) Co-IP analysis with Flag-UBE4B immunoprecipitates from lysates of 293T cells transfected with the indicated plasmids. Immunoblotting was performed with lysates using the indicated antibodies.

**Figure S2.**
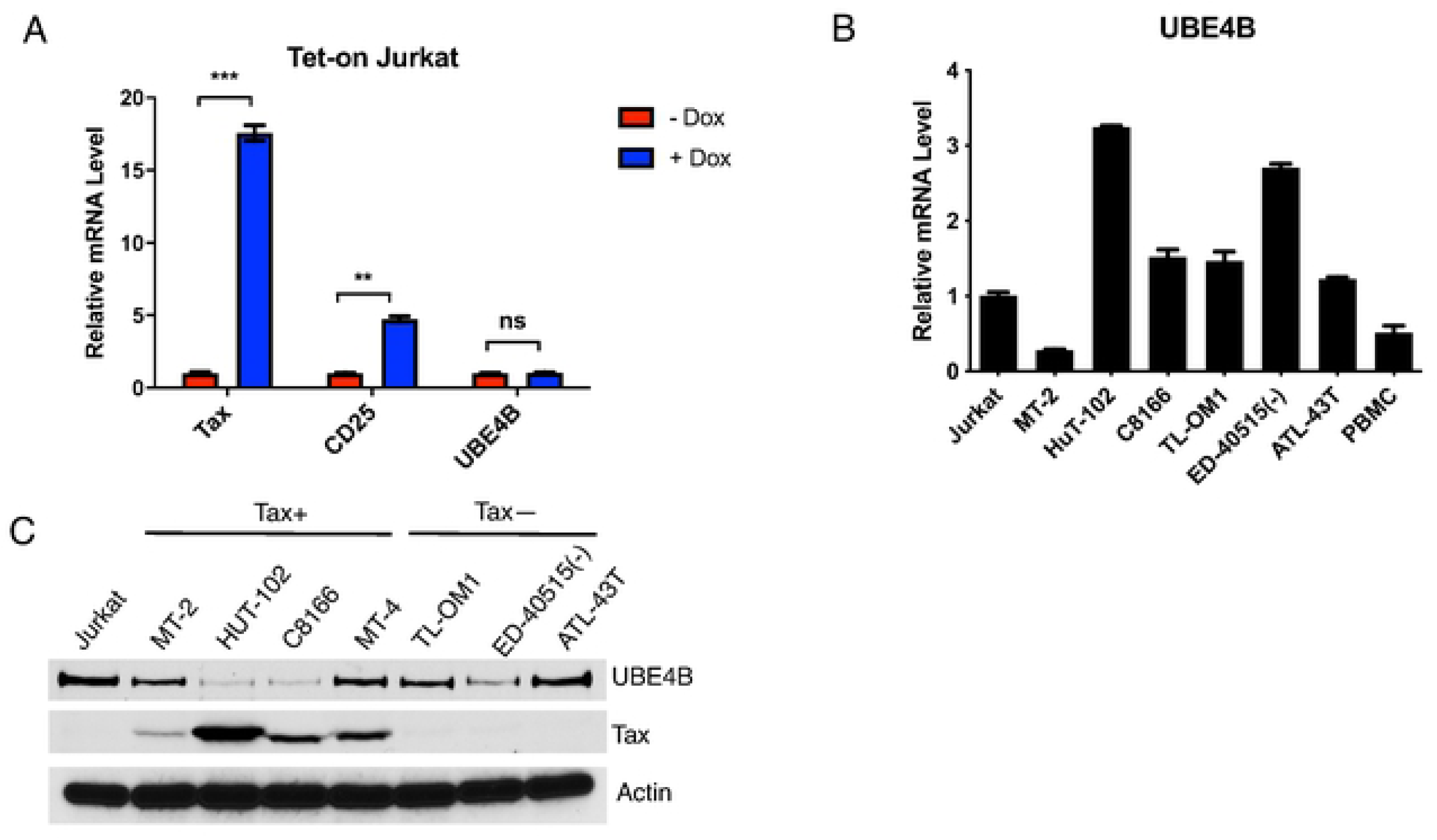
Tax does not upregulate the expression of UBE4B. (A) qRT-PCR of Tax, CD25 and UBE4B mRNAs in Jurkat Tax Tet-on cells treated either with doxycycline (Dox) or DMSO. (B) qRT-PCR of UBE4B mRNA in Jurkat, ATLL cell lines, and PBMCs. (C) Immunoblotting was performed with the indicated antibodies using whole cell lysates from Jurkat, Tax+ and Tax-ATLL cell lines.

**Figure S3.**
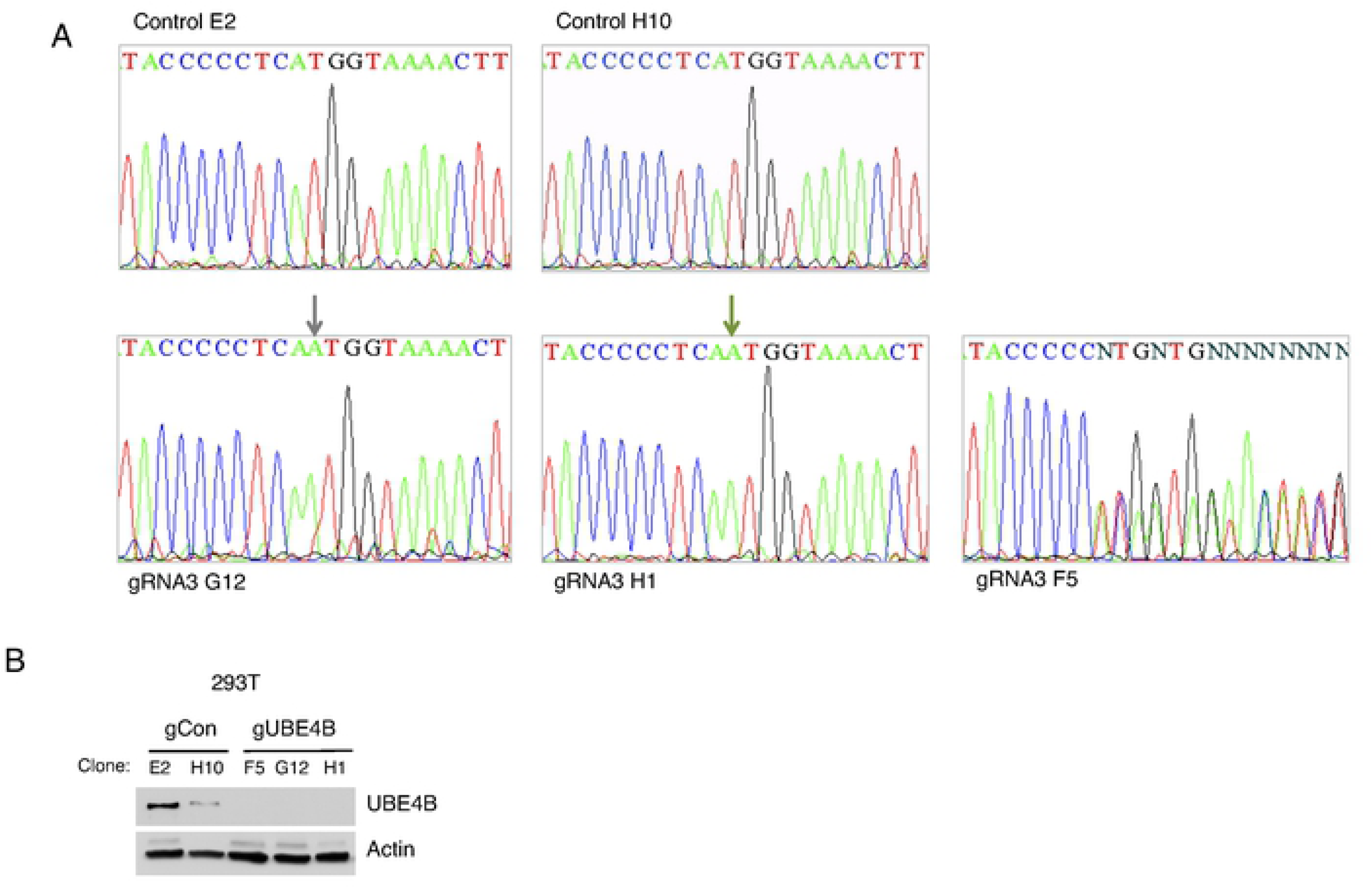
Characterization of UBE4B knockout 293T clones. (A) DNA sequencing chromatograms of PCR-amplified UBE4B exon 10 from genomic DNA derived from wild-type (E2, H10) and UBE4B KO (G12, H1, F5) 293T cell clones. UBE4B KO clones G12 and H1 both have an adenine insertion. (B) Immunoblotting was performed with the indicated antibodies using lysates from wild-type (E2, H10) and UBE4B KO (G12, H1, F5) 293T cell clones.

**Figure S4.**
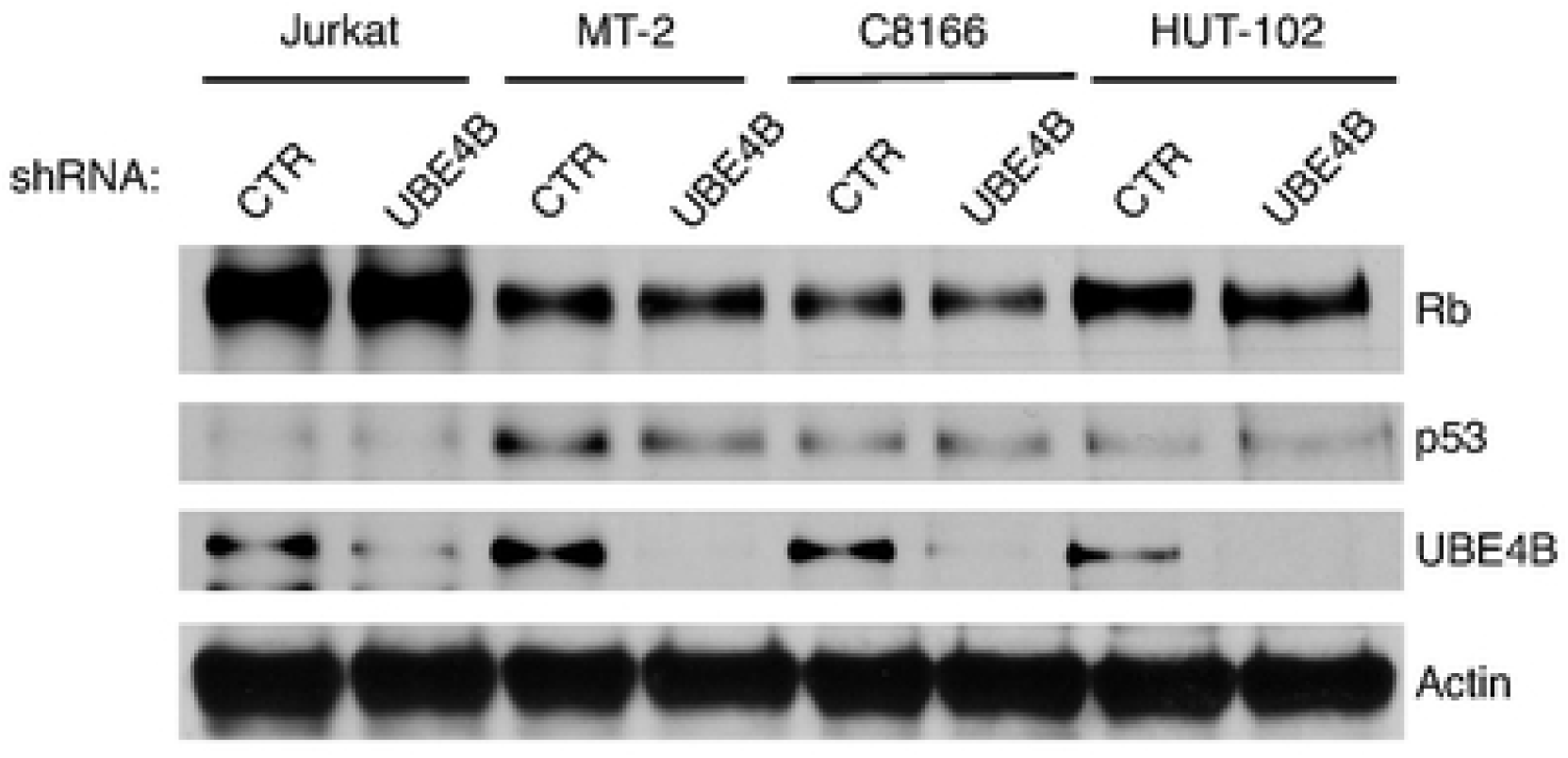
UBE4B does not promote Rb and p53 degradation in HTLV-1 transformed cell lines. Immunoblotting was performed with the indicated antibodies using lysates from Jurkat, MT-2, C8166 and HUT-102 cells expressing control or UBE4B shRNAs.

**Figure S5.**
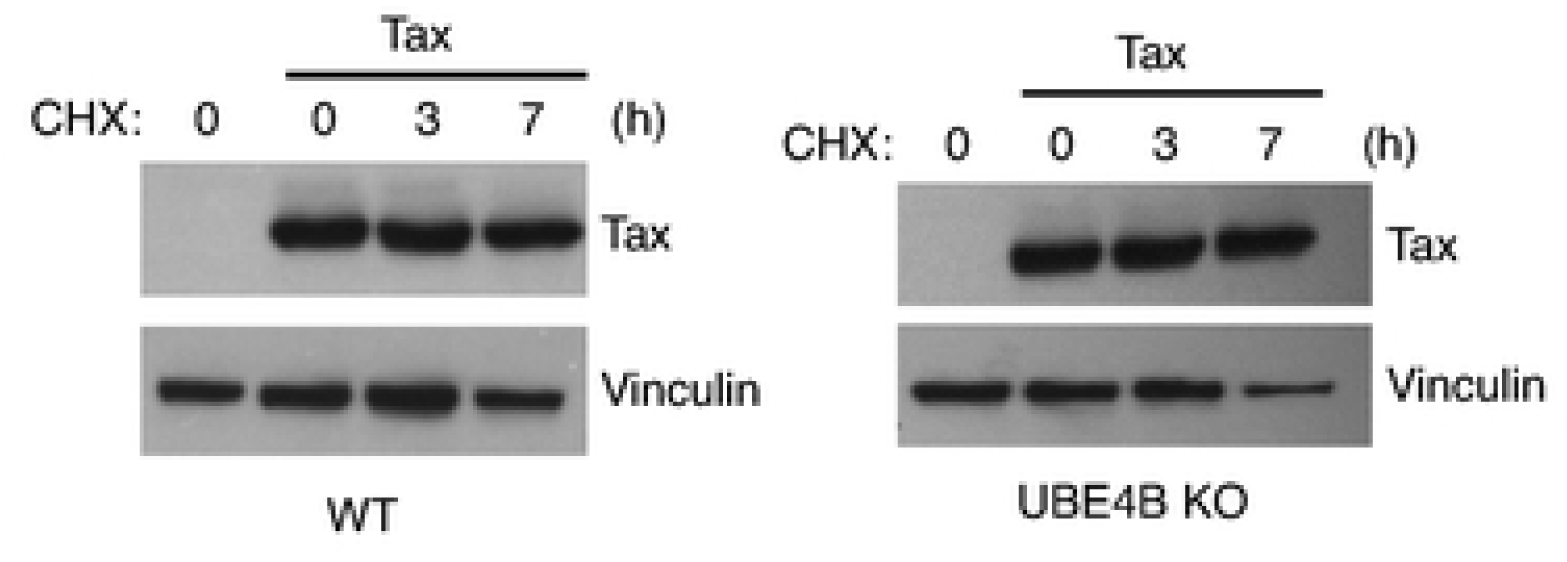
UBE4B does not destabilize Tax. Cycloheximide chase assay with lysates from wild-type and UBE4B KO 293T cells (clone H1) transfected with Tax and treated with cycloheximide for the indicated times. Immunoblotting was performed with the indicated antibodies.

